# Evaluation of Long-Short Term Memory Networks for M/EEG Source Imaging with Simulated and Real EEG Data

**DOI:** 10.1101/2022.04.13.488148

**Authors:** Lukas Hecker, Moritz Maschke, Rebekka Rupprecht, Ludger Tebartz van Elst, Jürgen Kornmeier

## Abstract

Magneto- and electroencephalography (M/EEG) are widespread techniques to measure neural activity *in-vivo* at a high temporal resolution but relatively low spatial resolution. Locating the sources underlying the M/EEG poses an inverse problem, which is itself ill-posed. In recent years, a new class of source imaging methods was developed based on artificial neural networks. We present a long-short term memory (LSTM) network to solve the M/EEG inverse problem. It integrates low computational cost, exploitation of both the coarse spatial but also the excellent temporal information from the EEG, input flexibility and robustness to noise. We compared the LSTM network with classical inverse solutions using both simulation data and real EEG data, recorded in epileptic patients during intracranial stimulation. The LSTM network shows higher accuracy on multiple metrics and for varying numbers of neural sources, compared to classical inverse solutions but also compared to our alternative architecture without integration of temporal information. The performance of the LSTM network regarding its robustness to noise and low localization errors renders it a promising inverse solution to be considered in future source localization studies and for clinical applications.

## 2 Introduction

Magneto- and electroencephalography (M/EEG) are non-invasive techniques to non-invasively measure human brain activity with high temporal resolution. The ease of use and excellent temporal resolution have made them one of the most used techniques in human neurophysiology. M/EEG suffers, however, from a considerably low spatial resolution. Inferring the active brain regions (henceforth called *sources*) given a signal measured on/at the scalp is not uniquely possible without adding further constraints, posing what is called the *inverse problem* (Grech et al., 2008; B. He, Sohrabpour, Brown, & Liu, 2018; Michel & Brunet, 2019; Nunez & Srinivasan, 2006; R. D. Pascual-Marqui, 1999). The reason for this is the underdetermined equation one has to solve: There are many more neurons (or ensembles thereof) in the brain than there are EEG electrodes. Furthermore, *volume conduction*, i.e., the summation of currents of various sources in the brain, also renders signals ambiguous and thus lowers the spatial resolution of the EEG further. Solving the inverse problem is useful for various reasons, e.g., to localize EEG responses in basic research (Kornmeier, Friedel, Hecker, Schmidt, & Wittmann, 2019) or aberrant activity as in pre-surgical epilepsy diagnostics (Mégevand, Hamid, Dümpelmann, & Heers, 2019). Furthermore, the disentanglement of the signal sources can be an advantage for brain state decoding or neurofeedback applications (van Lutterveld et al., 2017).

In order to uniquely solve the inverse problem, one has to introduce some constraint on the solution. One popular constraint is the *minimum norm*, used in various ways in a group of inverse solutions referred to as Minimum Norm Estimates (MNE; Dale & Sereno, 1993; Hämäläinen & Ilmoniemi, 1994; Ioannides, Bolton, & Clarke, 1990). These inverse solutions aim to find the source configuration that minimizes the required power to generate a given potential at the scalp electrodes. Low Resolution Electromagnetic Tomography (LORETA) and its iterations are famous proponents of the MNE-family which have shown low localization errors (R. Pascual-Marqui, Michel, & Lehmann, 1994; R. D. Pascual-Marqui, 2007).

In recent years, more investigations were conducted on the incorporation of the rich temporal information of the M/EEG to calculate inverse solutions (Castanõ-Candamil et al., 2015; Dinh, Samuelsson, Hunold, Hämäläinen, & Khan, 2021; Gavit, Baillet, Mangin, Pescatore, & Garnero, 2001; Gramfort, Kowalski, & Hämäläinen, 2012; Gramfort, Strohmeier, Haueisen, Hämäläinen, & Kowalski, 2013), although this aspect seems to be often overlooked. Incorporation of the temporal dimension poses an advantage both in handling noisy measurements and in accumulating evidence for weaker sources (e.g., Baron, Duarte, Wakin, Sarvotham, & Baraniuk, 2009; Tropp, Gilbert, & Strauss, 2005).

In addition to analytical approaches, Artificial neural networks (ANNs) are an emerging tool to tackle inverse problems. ANNs have made their first appearance in 1943 and saw an important leap in development with the development of the perceptron (McCulloch & Pitts, 1943; Rosenblatt, 1958). A couple of decades later, ANNs found their first application in solving the inverse problem of the EEG based on single and few dipoles (e.g., Abeyratne, Zhang, & Saratchandran, 2001; Robert, Gaudy, & Limoge, 2002). With the large increase of computing power and data resources in the past two decades, ANNs have gained in popularity and are now used successfully in a variety of tasks, e.g., image classification (Krizhevsky, Sutskever, & Hinton, 2012) and classification of single trial EEG (Schirrmeister et al., 2017). With this leap in technology, ANNs are now being reconsidered to solve the inverse problem in M/EEG and various research groups are starting to develop and refine architectures (see Awan, Saleem, & Kiran, 2019; Fedorov, Koshev, & Dylov, 2020; Razorenova et al., 2020; Zorzos, Kakkos, Ventouras, & Matsopoulos, 2021 for reviews). Deep ANNs were considered for inverse problems in other domains, too (Jin, McCann, Froustey, & Unser, 2017).

In our recent study, we proposed a simple convolutional neural network (CNN) termed ConvDip (Hecker, Rupprecht, van Elst, & Kornmeier, 2020) that solves the EEG inverse problem on single time points. As input, ConvDip takes a single time point of EEG and outputs the corresponding estimated source vector, thereby creating distributed inverse solutions independent of temporal context. Pantazis and Adler (2021) developed two different model architectures. The MLP model was constructed similarly to ConvDip, the difference being that there was no first convolutional layer and an alternative output structure. The output vector of the last layer only contained the position (x-, y- and z-coordinate) of one, two or three sources. Thereby, the number of expected sources must be known in advance. Despite promising results and their efficient design, these models discard a considerable amount of information by ignoring the temporal dimension and reduce the spatial pattern to merely a predefined number of dipoles.

In the first paper on ANN-based inverse solutions within the last decade, Cui et al. (2019) introduced a long short-term memory (LSTM) network trained on simulated EEG data which was designed to find the location and time-course of single-dipole sources. Since the model transforms one spatiotemporal input into a spatiotemporal output of the same length, it is considered a sequence-to-sequence (seq2seq) model. This was an important idea, resulting in a leap in ANN-based inverse solutions, albeit the simulation framework served rather as a proof-of-concept.

Sun, Sohrabpour, Ye, and He (2020) followed a different approach and designed a spatiotemporal CNN called *SIFNet*. The model transforms an EEG sequence of fixed length (500 samples) to an output of summed source activity. Unlike the architecture of Cui et al. (2019), a sequence-to-one (seq2one) model was implemented. Albeit limited by the lack of a temporal dimension of the output vector, the model showed promising results compared to standardized LORETA (sLORETA). Particularly, it showed great robustness to varying levels of noise in the input data, which appears to be one key advantage of ANN-based inverse solutions over classical approaches. Furthermore, in scenarios where the temporal dimension of source activity can be discarded (e.g., when localizing a segment of epileptiform activity) this approach may prove especially useful. The model was improved and data simulations were carried out using neural mass models Sun, Sohrabpour, Worrell, and He (2022).

Another spatiotemporal model developed by Pantazis and Adler (2021) harnesses temporal context using two-dimensional spatiotemporal filters and four fully-connected layers. Similar to the author’s earlier MLP architecture described above, the output vector represented dipole positions. Therefore, this architecture also yields seq2one transformations.

An interesting approach was published by Huang et al. (2020), who designed the Data-Synthesis-Based Denoising Autoencoder (DST-DAE). The architecture is based on an autoencoder network with convolutional (encoding) and deconvolutional (decoding) layers. The architecture requires a fixed-size temporal dimension, which was set to 40 time points. Although the DST-DAE was trained with a single and two source patches, the model showed promising results to reliably estimate both source positions and their sizes at varying levels of noise.

Given the high temporal resolution of the M/EEG in the range of 1 millisecond (**?**) and estimates about the durations of neural processing steps in the range between 60 ms and 120 ms (Michel & Koenig, 2018), the temporal dimension (i.e., EEG data from neighboring time points as input) can contain important information for source localization. One approach to add memory to any inverse solution was proposed by Dinh et al. (2021), who incorporated temporal context into dynamic statistical parametric mapping (dSPM, Dale & Sereno, 1993) using an LSTM network. The great advantage of this approach is that the LSTM operates entirely data-driven without making assumptions on the temporal progression of source activity. One drawback of this method, as the authors claim, is that temporal context can only be used from past but not from subsequent samples. Furthermore, the model needs a few samples to build a sufficient history, which means that the first few time points do not benefit from any temporal context. This is reasonable for online source analysis where only previous time points are available. However, the method misses out on information during offline analysis, where both preceding and subsequent time points are available as temporal context.

The ANN-based inverse solutions for distributed dipole models all suffer from the high computational complexity given by the large number of dipoles (typically *≈* 10^3^*to*10^4^ vertices) that directly determine the size of the output layer, leading to large numbers of parameters. Jiao et al. (2022) proposed a *Graph Fourier Transform* of the vertices’ adjacency matrix to reduce the number of dipoles. This works by selecting only a subset of the eigenvectors required to represent sparse source configurations, which in their case led to a reduction from 2,052 to 615 output nodes, thereby minimizing the number of parameters and thus computation time of their proposed bidirectional LSTM architecture. A technique with the purpose of compressing the output vector in a biophysically meaningful way using edge sparsity was presented by Wei et al. (2021). The technique is based on the assumption that any EEG topography can be explained using a sparse subset of a confined set of basis functions.

Finally, Sun et al. (2022) trained an LSTM model using biophysically motivated simulated pairs of source and EEG data. Therefore, the authors applied neural mass models (Breakspear, 2017), thereby training the ANN with realistic patterns of neural activity. The proposed model architecture consisted of a spatial and temporal module. The spatial path consisted of three residual blocks (i.e., residual neural networks with skip connections, K. He, Zhang, Ren, & Sun, 2016). The temporal is attached to the spatial path, consisting of three LSTM layers. The output consisted of the spatio-temporal source estimate.

In order to bring ANN-based inverse solutions out of the infancy state there are some issues that need to be addressed. Each of the existing ANN-based inverse solvers has at least one of the following shortcomings: (1) Large number of parameters, (2) rigid source outputs (i.e., single/ few fixed dipole position/s), (3) inability to exploit temporal information flexibly.

In this work we present an extension of the initial ConvDip, a bidirectional LSTM network which solves the M/EEG inverse problem for distributed dipole models in a seq2seq manner. Inspired by Godard, Matzen, and Uyttendaele (2018), we created a list of criteria an ANN-based algorithm for solving the inverse problem should optimally fulfil:

1. **Accessibility:** The ANN should be computable on common computers with the number of parameters not exceeding 1,000,000.
2. **Flexible input:** The ANN should be able to work both on single time instances and time series data (e.g., event-related potentials) of varying length.
3. **Broadly applicable priors:** The assumptions on the number, size and shape of sources as well as assumptions on temporal structure of time courses should be chosen broadly to promote adaptive inverse solutions.
4. **Robustness to noise:** The ANN should be capable of handling signals with variable signal-to-noise ratios (SNRs) appropriately, especially when applied to online EEG data.

Note, that these criteria don’t just affect the design of the network but also the properties of the simulated training data. In order to fulfil criterion (1) we have tried to keep the number of trainable parameters as low as possible. Furthermore, we analyzed the sparsity-performance trade-off. In conjunction with criterion (2) we decided to use a sequence-to-sequence (seq2seq) LSTM architecture which accepts inputs of different numbers of time points (see Fig. 1 C). Criterion (3) is fulfilled if the simulations are maximally diverse. This regards parameters like the number of source patches, their amplitudes, sizes, shapes as well as their temporal progression. We aimed to define particularly large value ranges of these parameters in our simulations, which are explained in more detail in the chapter *Source and EEG Simulation* and outlined in Table 1. Criterion (4) is handled in conjunction with criterion (3) since the ANNs’ ability to correctly estimate sources from noisy signals to some degree depends on the noisiness of the data it is confronted with. We therefore simulated trials with a wide range of signal-to-noise ratios (SNRs). To our best knowledge, there is currently no ANN-based algorithm for the M/EEG inverse problem that satisfies all of these criteria.

**Table 1:** Simulation Parameters. Diameters are reported in neighborhood orders, the higher this number the larger the diameter.

| Parameter | Generative Model 1 | Generative Model 2 | Domain |
| --- | --- | --- | --- |
| Number of dipole positions | 1284 | 8196 | spatial |
| Number of sources | [1, 10] | [1, 15] | spatial |
| Diameter | [1, 2] | [1, 5] | spatial |
| Source time course frequency spectrum | $[1/f^0, 1/f^3]$ | $[1/f^0, 1/f^3]$ | temporal |
| EEG signal-to-noise ratio | [1, 100] | [0.1, 100] | temporal |

**Figure 1:**
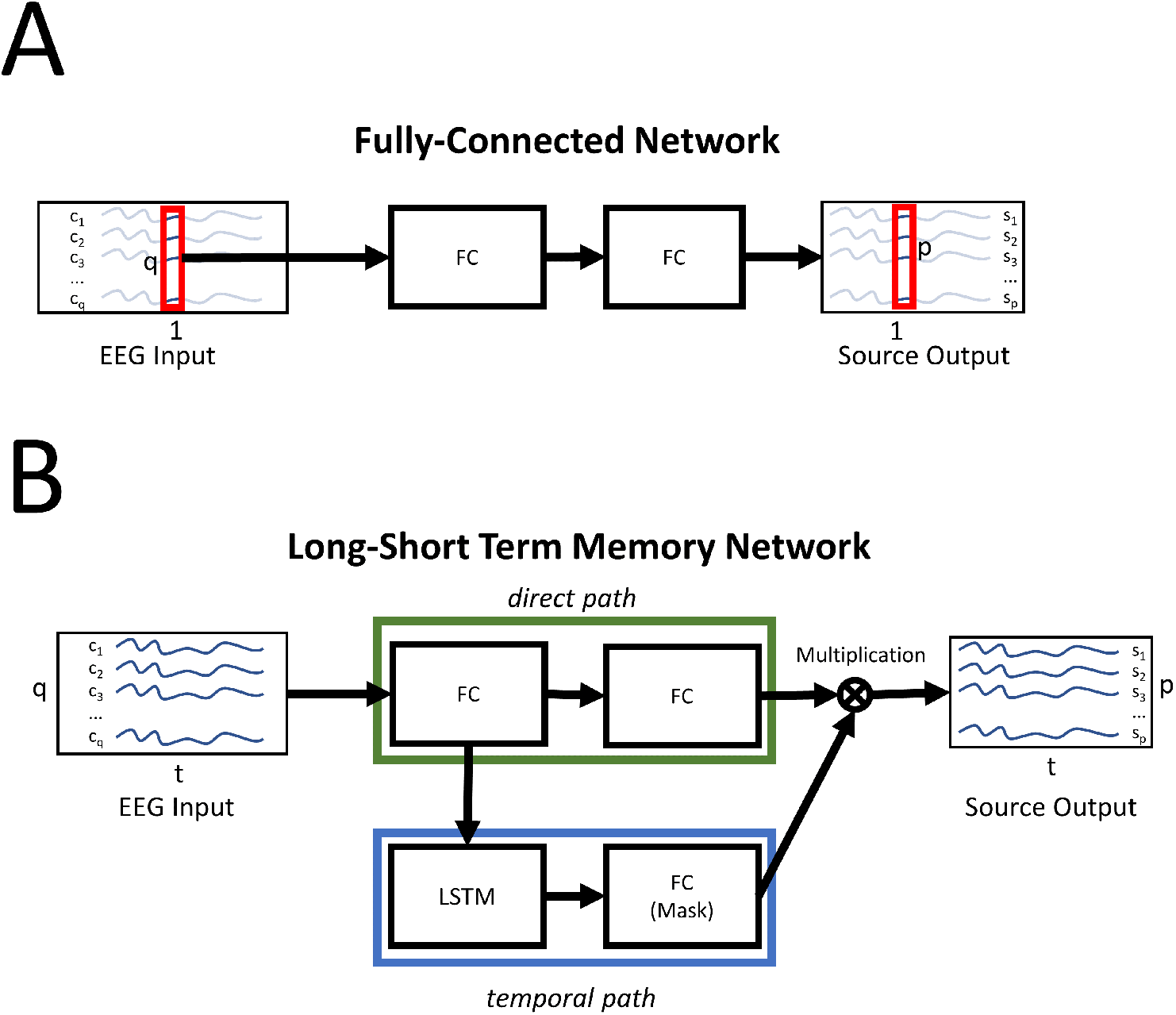
Neural network architectures. A: The fully-connected (FC) network with two hidden layers. B: The LSTM network. The green box indicates the direct path where the source is estimated without temporal information. The blue box indicates the temporal (memory) path, which creates a mask. The source estimate from the direct path and the mask of the temporal path are multiplied, which yields the final source estimate. q: number of electrodes, p: number of dipoles, t: time points, *c*_*i*_: channel at index *i, s*_*j*_ dipole at index *j*.

In order to compare the performance of the presented ANNs and classical source localization methods, we have designed the following twofold procedure: (1) We generate a new set of simulated source- and EEG-data pairs to serve as a ground truth. In order to avoid evaluating the models on the same type of data that they were trained on (*invere crime*) we have made several changes to the generative model, i.e., the forward model and simulation parameters. (2) In an additional evaluation step, we replaced the simulated evaluation data by real EEG recordings of intracranial electric simulation of the brain. For this, we used a data set which was published by Mikulan et al. (2020). These data constitute a more realistic scenario where real brain recordings are used, albeit the brain activity was induced externally.

## 3 Methods

### 3.1 Forward Model

We used an anatomical template brain “fsaverage” (Fischl, Sereno, Tootell, & Dale, 1999) by the Freesurfer image analysis suite^1^. EEG simulations were carried out using a precomputed three shell boundary element method (BEM; Fuchs, Kastner, Wagner, Hawes, & Ebersole, 2002) forward solution as provided by mne-python (v20.3, Gramfort, Luessi, et al., 2013). Each shell (brain, skull & scalp tissue) was composed of 5120 vertices. The conductivity was set to 0.3*S/m*^2^ for brain and scalp tissue, and 0.06*S/m*^2^ for the skull.

In order to spare computational resources, we chose a source model with low spatial resolution of *p* = 1, 284 dipoles with icosahedral spacing. For the EEG electrodes we used the Easycap-M1 layout consisting of *q* = 61 electrodes of the 10-20 system. Using the forward model and the parameters described, we calculated a leadfield *L ∈* ℝ ^*q×p*^, that assigns a gain value to each electrode for every source dipole in the brain. Therefore, it is often referred to as *gain matrix*. A fixed orientation of dipoles perpendicular to the cortical surface of the cortex was used (Michel & Brunet, 2019). This forward model was used to generate training data, wherefore we term it generative model 1 (GM-1).

We then defined a second forward model with slightly different parameters which was used to create the simulated source and EEG data for the *test set*, termed GM-2. The rationale behind this is to mitigate the problem of *inverse crime*, which states that results can be overoptimistic of the same generative model is used to create and test some model. The electric conductivity of the three shells was changed to 0.332*S/m*^2^ for the brain and scalp and to 0.0113*S/m*^2^ for the skull, thereby changing the skull-to-brain conductivity ratio from 1:50 to 1:25. The distribution of dipoles along the cortical surface was changed to a recursively subdivided octahedral spacing with *p* = 8, 196 dipoles, more than six times more dipoles than the model was trained on. Transformation of sources created with GM-2 to the source space of GM-1 was achieved using K-nearest neighbor interpolation with *K* = 5 neighbors. In this procedure, each dipole in the source space of GM-1 is assigned the mean activity value of the 5 nearest neighbors in the GM-2 source space.

### 3.2 Source and EEG Simulation

Our goal with the design of the simulation was to make as few assumptions as possible with respect to spatial and temporal properties. The parameters in our simulation are given in Table 1. For each simulated sample, the number of active sources was chosen randomly between 1 and 10. Furthermore, for each active source patch we randomly chose its diameter, maximum dipole moment and the frequency spectrum of its time course. Diameters were chosen in terms of neighborhood orders. A neighborhood order of 1 corresponds to single dipoles, whereas an order of 2 includes all direct neighbors of some dipole. The use of extended sources (as opposed to single dipoles) is motivated by numerous studies showing that neural sites of activity are (mostly) coherent in space and time (Destexhe, Contreras, & Steriade, 1999; Leopold, Murayama, & Logothetis, 2003).

Each simulated sample of source- and EEG activity contained 20 time points. The time course was chosen randomly at a frequency spectrum defined by 1*/f*^*β*^, whereas *β* was randomly chosen between 0 and 3 for each source individually. This approach yields highly variable time courses from aberrant (e.g., as in noisy raw EEG recordings) to smooth (e.g., as in event-related potentials). The source simulation yielded a source matrix *Y ∈* ℝ ^*p×t*^ with *p* = 1284 dipoles and *t* = *{*1, 2, …, 20*}* time points. This source matrix was multiplied with the leadfield, which yields the simulated and noise-free EEG *M ∈* ℝ^*q×t*^:

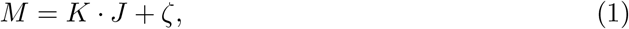

Noise of a white frequency spectrum was then added to the simulated EEG, scaled such that a randomly chosen SNR (within the range given in Tab. 1) was achieved. SNR was defined as the ratio of the averaged global field power (GFP) of signal and the averaged GFP of the noise.

Prior to training, each column of the EEG matrix was scaled by its Euclidean norm and each sample was scaled such that its absolute maximum was equal to one. Furthermore, common average referencing was applied to all EEG data.

### 3.3 Neural Networks

We will describe and test two different ANN architectures: (1) A fully-connected network (ANN-FC) similar to the on in our previous publication Hecker et al. (2020) and (2) the LSTM network (ANN-LSTM). The neural networks were built and trained using Python and the machine learning libraries Tensorflow 2.5.0 (Abadi et al., 2016) and Keras 2.5.0 (Chollet et al., 2015). We designed each ANN to have *≈* 1, 000, 000 parameters which we deem a good balance between computational complexity and model capacity. Mathematical notation of the three architectures are given in Appendix A.

#### 3.3.1 Fully-Connected Network

The FC network is a simple multilayer perceptron containing 2 hidden layers with *k* = 550 neurons each (Fig. 1, A) and an output layer of with neurons equal to the number of dipoles in the source model. ReLU activation functions were applied after each hidden layer. The total number of parameters sums up to 1, 046, 284. The model can be considered a simplified version of the previously published ConvDip model, which despite its successful performance involved unnecessary features Hecker et al., 2020. Specifically, we have removed the initial convolutional layer and have left out the 2D-interpolation of the EEG data, which led to better performance.

The FC network receives a single time point of EEG data of *y*_*t*_ *∈* ℝ^*q*^, with *q* channels. After the forward pass (see Appendix A for a mathematical description thereof) it outputs the inverse solution *x ∈* ℝ^*p*^ with *p* dipoles.

#### 3.3.2 Long Short-Term Memory Network

The LSTM network has a more complex architecture since its computing units are not simple neurons but so-called LSTM cells. These cells allow the network to retain (and forget) information from previous time steps and thus foster predictions that make use of the temporal context. This sets the model apart from the FC network described before. An LSTM cell consists of four different components: The input gate, forget gate, cell state and output gate. Multiple LSTM cells can comprise one LSTM layer with the advantage of capturing diverse temporal patterns in the data. The forward-pass of an LSTM network is described in detail in Appendix A. In an effort to facilitate the incorporation of the whole temporal context we have used an implementation of the bidirectional LSTM, which can not only utilize past time points but also subsequent time points (Schuster & Paliwal, 1997). Our LSTM architecture follows a similar approach as Dinh et al. (2021) with a direct non-temporal sub-network consisting of two FC layers ‘a 230 and 1284 neurons. A second path branch which we call contextualization path is attached to the first fully-connected layer and consists of a single LSTM consisting of 256 cells (128 per direction) and is followed by a large fully-connected layer of 1284 neurons, which is termed the mask. This layer was decorated with a sigmoid activation function and serves as a filter which sets dipole locations that have only little evidence to be active to zero. The mask layer is finally multiplied with the second fully-connected layer of the direct path, resulting in a temporally contextualized source estimate. The total number of parameters in this LSTM network is 1, 009, 158.

The input of the LSTM network is a 2D EEG *Y ∈* ℝ^*q×t*^ with *q* channels and *t* time points. The number of channels is a fixed parameter, whereas the number of time points (*t*) can vary. The output of the network is the inverse solution *X ∈* ℝ^*p×t*^ with *p* dipoles and *t* time points (see Appendix A for a mathematical description of the LSTM’s forward pass).

#### 3.3.3 Training Procedure

We have implemented a data generator that generates new pairs (samples) of source and EEG activity during training. This data generator returned a batch of 2,568 new samples (i.e., pairs of simulated source- and EEG-data) every epoch. The ANNs were trained on the same batch for 30 times before a new set of samples was generated. The ANNs were trained for 30,000 epochs, resulting in a total 2,568,000 samples. A validation set was created independently of the training data and was used to monitor generalization during training and to restore the best set of weights based on the validation loss after all epochs were finished.

The loss was calculated by *cosine-similarity* between predicted and target source vectors, a distance metric that penalizes errors in source patterns but not in absolute source amplitudes. As an optimizer we chose adaptive momentum estimation (ADAM, Kingma & Ba, 2014) with a learning rate of 0.001.

Due to the scaling of the training data and the scale-agnostic loss function, it follows that the ANNs will predict sources that do not reflect the original scaling. We rescaled the predicted source such that the difference between the measured EEG (*M*) and the EEG produced by the predicted source (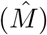) was minimally correlated with the measured EEG:

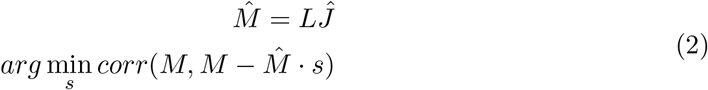

where L is the leadfield matrix *L ∈* ℝ^*q×p*^ and 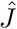 is the source estimate from the ANNs output layer.

### 3.4 Evaluation

The evaluation of the ANN’s inverse solutions and their comparison with other inverse solutions is realized in two steps: (1) Using simulated pairs of source- and EEG-data and (2) based on real data.

#### 3.4.1 Evaluation with Simulated Data

In order to avoid what is called the *inverse crime* we created a new forward model with some modified parameters and changed the simulation parameters. We refer to this alternative model as *Generative Model 2* (GM-2). In order to evaluate the ANNs with data outside their known range, simulations were carried out using slightly more challenging parameter ranges (Tab. 1). Using these parameters, we simulated 5,000 pairs of source- and EEG-activity in order to assess the accuracy of all solvers.

The following metrics were used where applicable to assess the accuracy of the inverse solutions compared to the ground truth:

- **Area under the ROC curve**. We calculated the area under the receiver operating characteristic curve (ROC-AUC) as described by Grova et al. (2006) and in our earlier work (Hecker et al., 2020). *AUC*_*far*_ is a metric that captures how well an inverse solution (1) finds the sources at the correct locations and (2) avoids false positives. *AUC*_*close*_ also captures (1) how well sources are correctly localized but also (3) how well the size was estimated. Therefore, *AUC*_*close*_ captures the dispersion of an inverse solution.
- **Mean localization error**. The Euclidean distance between the locations of the predicted source maximum and the target source maximum is a common measure of inverse solution accuracy as it captures the ability to localize the source center accurately. MLE was calculated between the positions of the maxima in the predicted source 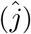 and the ground truth source (*j*). This metric is only suitable for calculating MLE when a single source patch is present.
- For multiple sources, we adapted the following procedure. First, we identified local maxima in both the true source vector *j* and the estimated source vector 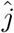. A voxel constituted a maximum if its value was larger than all of its neighboring voxels. This yielded many local maxima, which had to be filtered in order to achieve reasonable results. Filtering involved the elimination of all maxima whose neighbors were not sufficiently active (*<* 10% of the maximum). This takes care of false positive maxima that do not constitute an active cluster of voxels. Then, we removed those maxima that had a larger maximum voxel within a radius of 30 mm of its own position. These procedures result in a list of coordinates of maxima for both *j* and 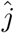. We then calculated the pairwise Euclidean distances between the two lists of coordinates of maxima. For each true source, we identified the closest estimated source and calculated the MLE by averaging these minimum distances. We further labelled those estimated sources that were *≥* 30*mm* away from the next true maximum as ghost sources. True maxima that did have an estimated source within a radius of 30 mm were labelled as found sources, whereas those true maxima that did not have an estimated maximum within a radius of 30 mm were labelled as missed sources. Finally, we calculated the percentage of found sources, i.e., the ratio of the number of correctly identified sources and the number of all true sources.
- **Normalized Mean Squared Error**. Calculation of the normalized MSE (nMSE) was done by normalizing both the true source vector *j* and the predicted source vector 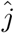 before calculating the MSE. By normalizing the vectors, we get rid of possible offsets and capture the overall differences in patterns more closely.

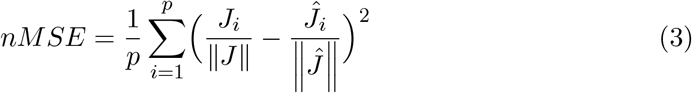

#### 3.4.2 Application to Real Data

In order to assess the accuracy of the LSTM in a more realistic setting with data from real brains we applied it to EEG data of intracranial electric brain stimulation of epileptic patients. The data used was published and reported by Mikulan et al. (2020). This open-source data set contains data of 7 patients undergoing pre-surgical epilepsy diagnostic. Each participant received single pulse electric stimulation (SPES) by intracranial electrodes at multiple locations. EEG was recorded during the stimulation and each location was stimulated multiple times.

The EEG was recorded at a sampling rate of 8,000 Hz at 256 electrodes on the equidistant Geodesic Sensor Net, HydroCel CleanLeads. The data has been cleaned by visual inspection by the data owners (e.g., removal of bad channels, artifacts, and bad epochs). Only the processed epochs (cropped from -250 ms to 10 ms) were provided. For preprocessing, the data was highpass filtered at 0.1 Hz cut-off (FIR filter; zero phase, Hamming window; automatic selection of length and bandwidth). For two patients (sub-05 and sub-07), a notch filter was applied at 50, 100, 150 and 200 Hz (FIR filter; zero phase; Hamming window; bandwidth = 0.1 and automatic length selection) to remove line noise and its upper harmonics. The data set owners generated epochs/trials from 300 ms prior to stimulation onset to 50 ms after. The baseline was corrected by subtracting the mean response between 300 ms and 50 ms before stimulation onset. Each participant was stimulated at between 7 - 14 different locations, resulting in a total of 55 runs. Note, that each run contained multiple repetitions of the same SPES. We calculated the averaged response (across repetitions) per patient and run. Next, we calculated the signal-to-noise ratio (SNR) for each run by dividing the absolute maximum value of the run’s global field power (GFP) by the average GFP during the baseline window. Finally, the trial was cropped to remove the peri-stimulation data segments, leaving only the EEG of the averaged response to the biphasic stimulation.

T1-weighted MR images with the faces masked by a low-resolution mesh for each patient were provided with the data set. Segmentation of the T1-weighted images was done using *freesurfer*. This resulted in an error in one subject (*sub-02*), wherefore we had to exclude all the related data. After segmentation of brain, skull and skin tissue we calculated the boundary element method (BEM) model with 5120 vertices in each of the three layers. Conductivity was set to 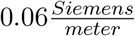 for the skull and to 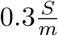 for the brain and scalp tissues.

The dataset contained digitized electrode positions which were co-registered with the MRI. The source space was set up using subdivided octahedrons, which resulted in 8196 dipoles in total. Finally, dipole orientations were assumed to be perpendicular to the surface of the cortex in order to reduce computational load.

As described above, the SPES data set contains EEG data from real brains stimulated with sinusoidal pulses. Figure 2 shows an exemplary average of 38 pulses. The ANNs described so far were trained with a diverse set of sources and EEG-data samples, containing single or multiple sources with varying spatial extent. All training data were simulated based on current knowledge about the typical source activity pattern generated by a real brain. The SPES data used in the current study contain electric pulses generated by implanted electrodes. The shape and temporal extent of this artificially generated signal may deviate from our assumptions about a typical brain-generated signal. In order to optimize our ANNs also for such patterns of activity, we can train it with samples that are closer to the SPES conditions, i.e., with single dipole sources matching shape and temporal extent of the artificial signals. We have therefore trained each ANN architecture (ANN-LSTM and ANN-FC) with the original training data (“Diverse”) and with SPES-adapted training data (“Single”) containing only a single source per sample.

**Figure 2:**
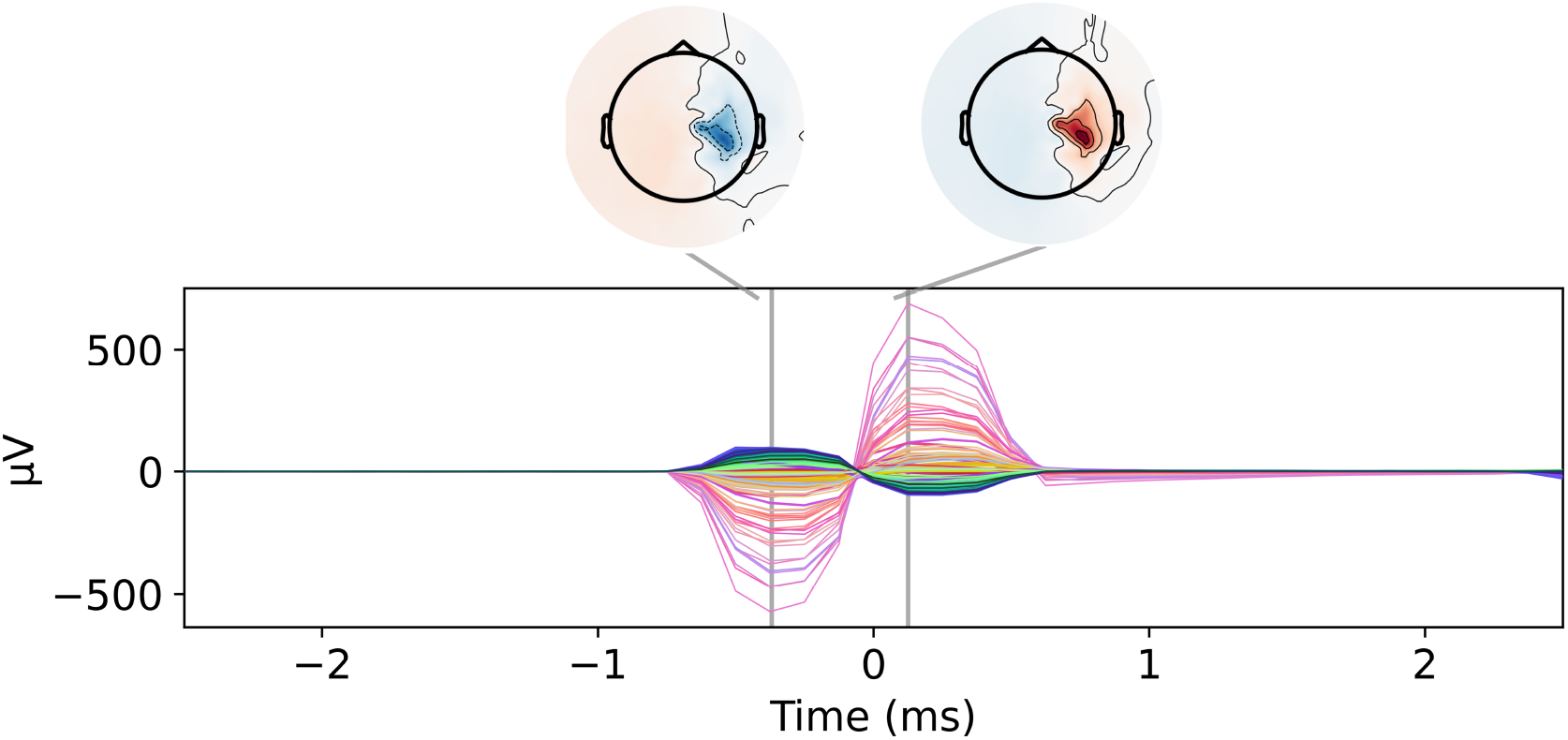
SPES-evoked potentials. Exemplary averaged response of 38 single-pulse electric stimulations (SPES) of a single patient. Note the short duration, high amplitude and sinusoidal shape of the EEG response.

For the evaluation of all solvers we calculated the Euclidean distance between the location of the stimulation electrode and the calculated position of the dipole that exhibited the maximum magnitude during the whole pulse. Other metrics could not be calculated since only the location of the stimulation electrodes and no spatial extent of the resulting cortical activity was known as a ground truth.

### 3.5 Inverse Solutions for Comparison

We compared our proposed network-based source analysis approaches (ANN-LSTM & ANN-FC) to a diverse set of commonly used approaches: exact (e)LORETA, dynamic Statistical Parametric Mapping (dSPM), Truncated and Recursively Applied Multi Signal Classification (TRAP-MUSIC), Linearly Constrained Minimum Variance (LCMV) Beamforming and

EPIFOCUS (Dale, Fischl, & Sereno, 1999; de Peralta Menendez, Andino, Lantz, Michel, & Landis, 2001; Mäkelä, Stenroos, Sarvas, & Ilmoniemi, 2018; R. D. Pascual-Marqui, 2007; Van Veen, van Drongelen, Yuchtman, & Suzuki, 1997). The code for these methods was developed in-house and is available as a free open-source python library ^2^. Regularization parameters were chosen optimally for each individual sample using generalized cross-validation (GCV) with 24 different magnitudes of regularization. Due to this careful procedure we did not whiten the data with a noise covariance matrix.

## 4 Results

### 4.1 Evaluation Based on Simulated Data

In this section shows the accuracy of the LSTM network compared to other solvers using 5,000 simulated samples of source and EEG activity as previously described. The simulations were generated using the *Generative Model 2*. Note, that this model contains an altered forward model with different dipole spacing and a broader set of stimulation parameters (see Tab. 1) compared to the parameters used during training.

Figure 3 and Table 2 depict the accuracy of the solvers concerning mean localization error (MLE), normalized mean squared error (nMSE) and area under the receiver operator curve (AUC). For single sources and source patches, the LSTM network produces significantly lower MLEs than all other solvers. The LSTM network produces 42 % lower MLE than the next-best solver (TRAP-MUSIC, *t* = 8.03, *p* = 3.74 *·* 10^*−*14^, *d* = 0.58). The AUC, a measure of global similarity, was highest with the ANN-FC, although this margin was not significantly different from the LSTM network (*t* = 1.63, *p* = 0.10, *d* = 0.10). The LSTM network outperformed all other solvers concerning nMSE by considerable margins. Compared to TRAP-MUSIC, which is closest to the LSTM’s performance, there is still a medium-sized improvement (*t* = 6.30, *p* = 1.33 *·* 10^*−*9^, *d* = 0.32).

**Table 2:**
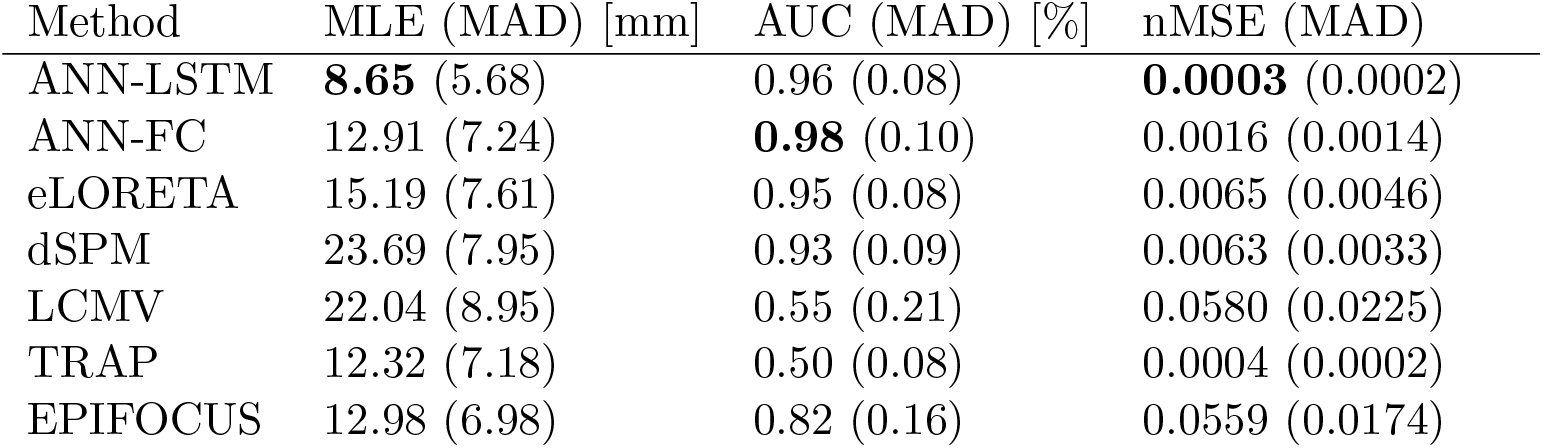
Evaluation Metrics for Single Sources. Medians and median absolute deviations (MAD) of the mean localization error (MLE), area under the ROC curve (AUC) and normalized mean squared error (nMSE). The best performing inverse solutions are marked in bold font.

**Figure 3:**
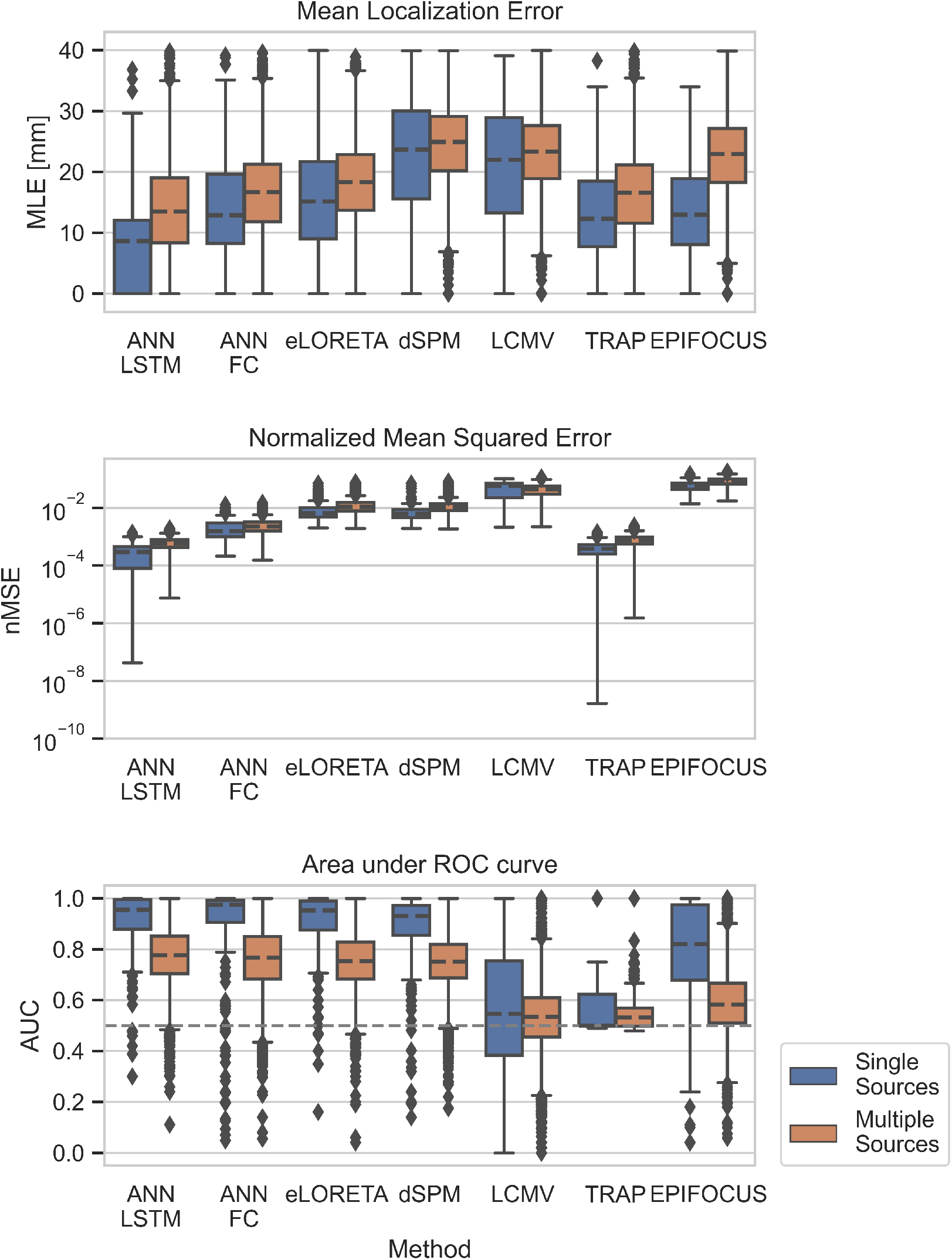
Accuracy Evaluation of all Solvers. Left: Performance on samples containing a single source or source patch. Right: Performance on samples containing between 1 and 15 sources or source patches.

The LSTM network’s performance is more distinguished when multiple sources are active at the same time (Fig. 3, right column). It achieves better accuracy on all observed metrics (Table 3). TRAP-MUSIC achieved the second-lowest errors concerning MLE and nMSE. However, both were significantly higher than the errors produced by the LSTM network (*t >* 23.70, *p <* 1.62 *·* 10^*−*117^, *d >* 0.40). AUC was highly similar among the ANN-based solvers and the minimum-norm-based solvers (eLORETA & dSPM), although the LSTM slightly outperformed its closest competitor (FC) by a significant margin (*t* = 9.01, *p* = 2.87 *·* 10^*−*19^, *d* = 0.10).

**Table 3:** Evaluation Metrics for Multiple Sources. Medians median absolute deviations (MAD) of the mean localization error (MLE), area under the ROC curve (AUC) and normalized mean squared error (nMSE). The best performing inverse solutions are marked in bold font.

| Method | MLE (MAD) [mm] | AUC (MAD) [%] | nMSE (MAD) |
| --- | --- | --- | --- |
| ANN-LSTM | **13.08** (6.33) | **0.78** (0.09) | **0.0006** (0.0002) |
| ANN-FC | 16.54 (5.75) | 0.77 (0.11) | 0.0022 (0.0012) |
| eLORETA | 18.22 (5.59) | 0.76 (0.09) | 0.0108 (0.0051) |
| dSPM | 24.88 (5.47) | 0.76 (0.09) | 0.0106 (0.0042) |
| LCMV | 23.33 (5.46) | 0.54 (0.10) | 0.0443 (0.0158) |
| TRAP | 16.41 (5.77) | 0.53 (0.04) | 0.0007 (0.0003) |
| EPIFOCUS | 22.64 (5.73) | 0.59 (0.11) | 0.0823 (0.0199) |

In summary, we find that the LSTM network achieved similar or (in most cases) significantly higher accuracy compared to all other solvers. This was particularly the case with MLE and nMSE and more pronounced when multiple neural sources were active simultaneously.

Figure 4 depicts the accuracy of all solvers with varying levels of sparsity (top row) and noise (bottom row). We find that TRAP-MUSIC is especially robust when SNR is considerably low (*<* 2). For SNRs above 2, however, the LSTM network achieves lower MLE and nMSE than all other solvers. For very sparse samples (1-15 active dipoles), we find that the advantage of the LSTM network over the FC network is greater compared to less sparse samples (50-100 active dipoles). This indicates that the LSTM network uses information in the temporal domain that allows it to produce sparser inverse solutions when necessary.

**Figure 4:**
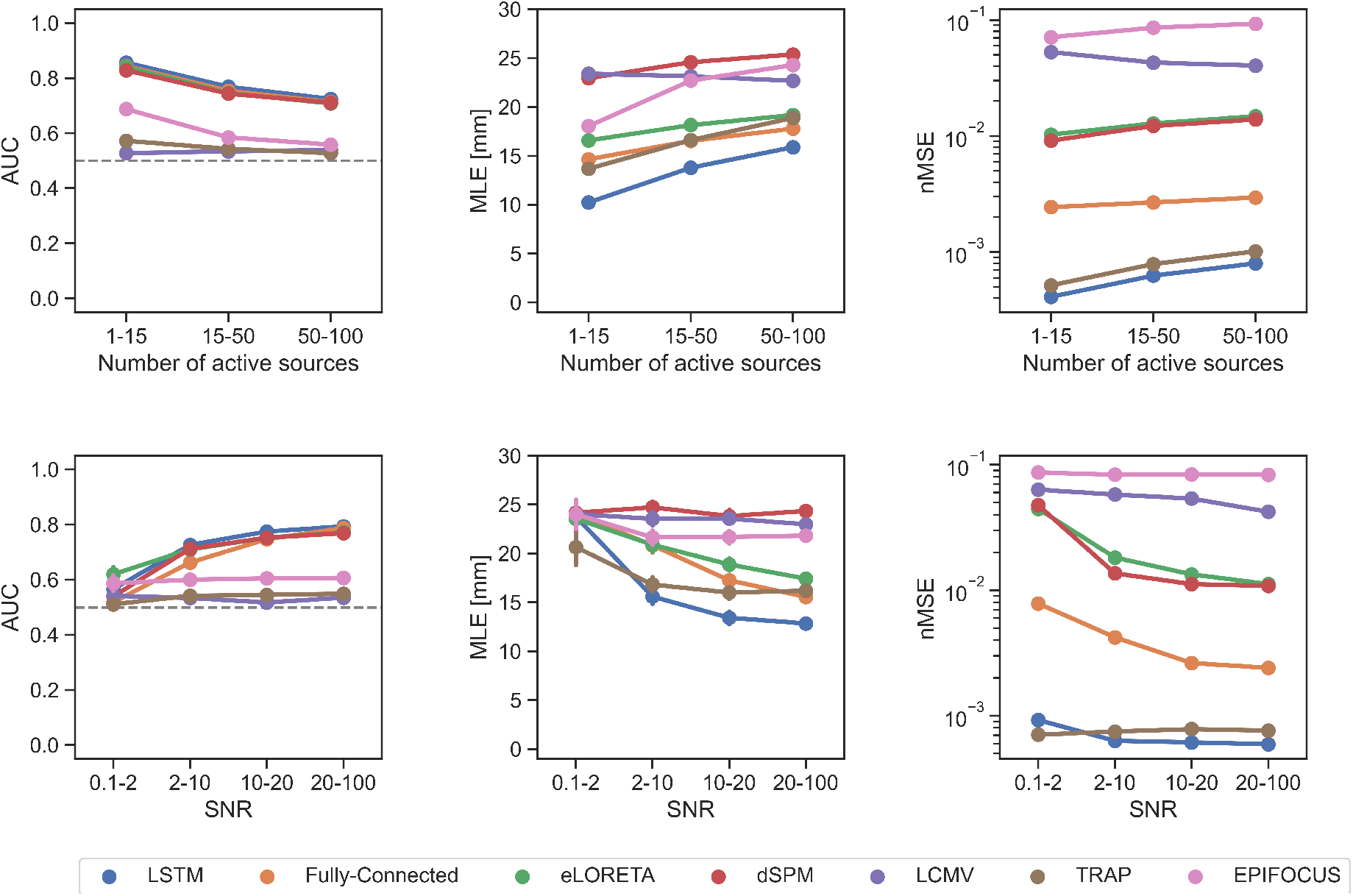
Accuracy with Varying Levels of Noise and Sparsity. Left column: Area under the receiver-operator curve (AUC). Center column: Mean localization error (MLE). Right column: Normalized mean squared error (nMSE).

### 4.2 Evaluation on Intracranial Stimulation Data

The following analysis shows the localization performance of all solvers on the intracranial SPES data set. The data of all patients and runs were pooled together for an overview and are depicted in Fig. 5. Each data point reflects the localization error between the position of the stimulation electrode (*ground truth*) and the dipole with maximum amplitude of the inverse solution produced by the respective solver. For the purpose of this evaluation we had to train individual ANNs for each patient. Particularly, we trained two separate ANNs per patient: (1) One ANN with a diverse set of simulation data (“Diverse”) and (2) a second network with simulations containing only single sources and source patches (“Single”) adjusted to the SPES data (see Application to Real Data).

**Figure 5:**
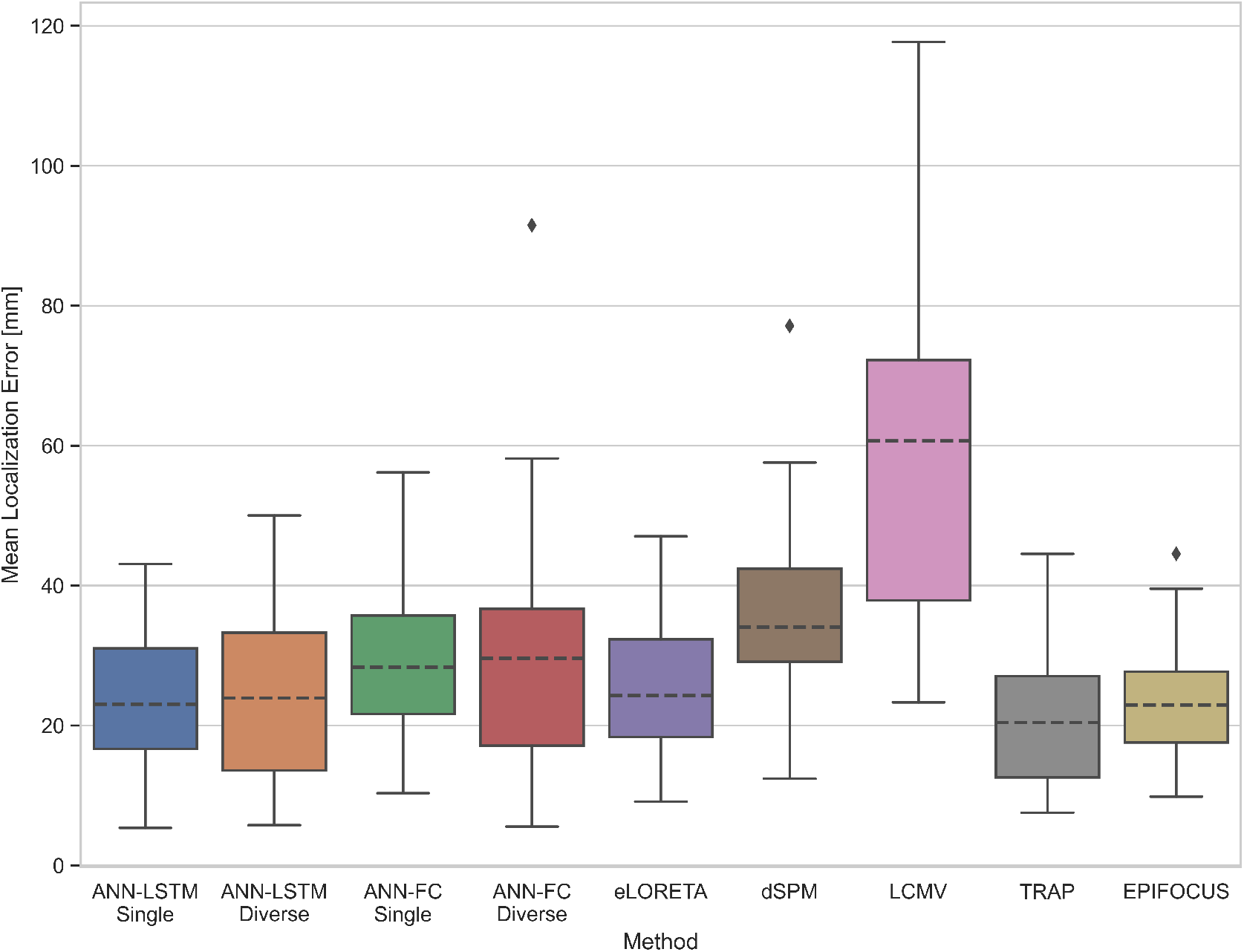
Localization Errors with Intracranial Stimulation Data.

TRAP-MUSIC achieved the lowest mean localization errors overall (*Median* = 20.5*mm*). The difference in MLE compared to the FC networks and ANN-FC Diverse) was statistically significant (ANN-FC Single: *t* = 3.81, *p* = 0.0003, *d* = 0.75, ANN-FC Diverse: *t* = 3.09, *p* = 0.003, *d* = 0.61), but not compared to the LSTM networks (ANN-LSTM Single: *t* = 1.44, *p* = 0.15, *d* = 0.19, ANN-LSTM Diverse: *t* = 1.37, *p* = 0.18, *d* = 0.22).

In Figure 5, it is obvious that the LSTM networks both yield lower MLE than the FC networks. The improvement of MLE between the *ANN-LSTM Single* over *ANN-FC Single* was 22.9% (*t* = 2.62, *p* = 0.011, *d* = 0.56) and the difference between the *ANN-LSTM Diverse* over *ANN-FC Diverse* was 23.6% (*t* = 3.34, *p* = 0.002, *d* = 0.42).

Interestingly, the ANNs that were trained only on samples containing single-sources (ANN-LSTM Single & ANN-FC Single) both achieved lower MLE compared to the counterparts that were trained on a diverse set of simulations. This difference was, however, not significant in both cases (*t ≤* 0.21, *p >* 0.83, *d <* 0.04). These results indicate that in cases of precise expectations about the pattern of neural activity the ANN’s training parameters can be adjusted to model these expectations, alebit the improvements are no very large.

In summary, we have evaluated the LSTM network using simulated and real EEG data from intracrancial stimulation of epileptic patients. With the simulations data, the LSTM achieved in most cases significantly higher accuracy in source localization results than the other source analysis methods. The benefit was particularly large when multiple sources were active simultaneously. Comparing our network approaches with each other, we found that the LSTM network outperformed the FC network particularly when the ground truth source pattern was sparse, i.e. when only few dipoles were active at the same time. Using the SPES dataset we calculated localization errors in a semi-realistic setting involving EEG recordings on patients that were stimulated intracranially. This approach has the advantage that the test data result from real human brains and at the same time a precise ground truth is available. With these data, the LSTM network shows comparable results with classical methods that are particularly designed to find sparse sources. Moreover, LSTM network models, taking advantage of the temporal information of EEG data, outperformed the FC networks, focusing only on time points.

## 5 Discussion

The EEG and MEG are widely used techniques to measure neurophysiological signals *in vivo*. Increasing their spatial resolution by solving the inverse problem, i.e. finding the neural sources given the signals measured on the scalp, is of major interest both for basic research and in the clinical context (e.g., localization of seizure onset zones in epilepsy). ANN-based analysis methods are gaining attention in the past years and appear a viable option to solve the inverse problem of the M/EEG (Cui et al., 2019; Dinh, Samuelsson, Hunold, & Hämäläinen, 2019; Hecker et al., 2020; Huang et al., 2020; Pantazis & Adler, 2021; Sun et al., 2020; Tankelevich, 2019).

In order to bring ANN-based inverse solutions out of the infancy state some issues need to be tackled. Each of the existing ANN-based inverse solvers has at least one of the following shortcomings: (1) Large number of parameters, (2) limited source outputs dimension (i.e., a few fixed dipole positions), (3) inability to exploit temporal information flexibly. In the present study we introduced an LSTM network which offers to solve these problems. The LSTM network was evaluated alongside a simple fully-connected (FC) ANN architecture and multiple established non-network source imaging methods using simulated and real data.

We have demonstrated that the LSTM network was able to produce inverse solutions the with higher (in few cases equal) accuracy compared the all other solvers tested. The LSTM excelled the most when multiple sources were active at the same time, where it achieved 25 % lower MLE, 2.6 % higher AUC and 17 % lower nMSE compared to the next-best non-ANN solver. Comparing the two ANNs with each other, the incorporation of temporal information led to a decrease in localization error and enhanced sparsity over solutions produced by the FC network. Interestingly, while the FC network produced solutions comparable to the minimum-norm based solvers (eLORETA & dSPM), the LSTM network produced solutions that behaved more similar to TRAP-MUSIC. This can be observed in Fig. 4, in which the LSTM network has similar (albeit, significantly lower) strengths and weaknesses concerning MLE and nMSE, but considerably higher AUC compared to TRAP-MUSIC.

Moreover, the LSTM network showed to produce accurate inverse solutions with varying SNRs. In a global measure of accuracy, we find that the LSTM suffers less from low-SNR compared the FC network, which we attribute to the added temporal information.

In a second evaluation step, the LSTM network was applied to real EEG recordings of intracranial stimulation data in order to test its accuracy under realistic conditions. This task combines multiple challenges. First, the forward model computation based on individual T1 scans is prone to errors since several parameters (e.g., conductivity) need to be guessed. Second, the stimulation electrode was inserted at varying angles, most of which did not align to be perpendicular to the cortical surface. This increases the difficulty to find the correct source locations since our forward models assumed fixed dipole orientations. Finally, the “sources” generated by the intracranial stimulation are presumably artificial compared to the expected distributed patterns of normal endogenous brain-electric activity. Furthermore, the stimulation has unknown extent and can potentially elicit cascades of cortico-cortical evoked potentials that may be localized by some solvers (Matsumoto, Kunieda, & Nair, 2017; Matsumoto et al., 2004). Solvers that are designed to find singular dipoles (e.g., TRAP-MUSIC) are thus expected to yield the lowest localization errors, whereas distributed EEG inverse solution (e.g., the proposed ANNs or eLORETA) may be disturbed by the aforementioned factors.

As expected, we find that TRAP-MUSIC and EPIFOCUS produce quantitatively the lowest localization errors on the simulation data among all methods tested. The localization errors produced by TRAP-MUSIC were, however, not statistically smaller compared to those produced by both LSTM networks. Again, comparing the ANN-based approaches with each other, the LSTM networks on the other hand outperformed the FC networks, which shows that the improvements through incorporation of temporal context make a change in real-world applications.

Currently it is still necessary to keep the number of parameters low such that ANNs can be trained on typical hardware available to researchers and clinicians. The reason for this is the comparatively long training time, which may limit the use of these techniques to researchers and clinicians with either appropriate computational resources. This problem becomes systematically larger with an increasing number of network parameters. One temporary solution is the use of small source models (1,000 - 5,000 dipoles) to shorten training times. An alternative solution to this problem could be publicly available pre-trained models and data sets. However, since ANN-based inverse solutions only work with a given electrode/sensor layout, anatomy and source space, pre-trained models could only be used for generic cases (i.e., template brains and electrode layouts) or would require retraining with the individual electrode layout and source space by replacing the input and output layers. Another option would be to compress the output space as described by Jiao et al. (2022), leading to fewer neurons in the output layer.

Of course, there are cases in which larger computational efforts could be justified, e.g., during presurgical procedures in epilepsy. In this case it may be reasonable to build ANNs with a larger number of parameters and to train it for longer on the appropriate hardware. In summary, we show that the ANN-based inverse solutions are in the vast majority of cases at an advantage over classical source analysis methods. Furthermore, we showed that the LSTM network makes use of the temporal information in the EEG, which leads to low localization errors, higher sparsity and more robustness to noise. As a proof-of-principle, we finally demonstrated, that the LSTM network, trained with simulated data, have an advantage over the FC network and other distributed dipole solutions (e.g., eLORETA, dSPM).

This work has thus demonstrated in various ways that ANNs, especially LSTM networks, are suitable to produce accurate inverse solutions in simulations, that are most often superior to classical non-network methods. In a second step we demonstrated that this good performance also and that this transfers to real-life settings. We are convinced that these methods can be used in basic research and by clinicians to produce more accurate inverse solutions and improve both our understanding of the brain and the outcome of surgical interventions in epilepsy.

Further optimization of these methods may be reached in developing physiologically plausible models of meso-scale brain-electric activity that allows for a fast generation of training data (see Sun et al., 2022 for an example). Furthermore, compression of the output layer of ANNs should be investigated further as a means to drastically reduce the number of parameters needed in the model.

Moreover, the remarkable similarity of the ANN-based results from the single source real brain data with the results from the classical approaches that optimized for such scenarios is further confirmation for this novel source analysis approach and renders it a viable choice, be it applications in epileptology, neurofeedback or ERP studies.

## 6 Data Availability

The simulations and neural network models that were presented in this work can be recreated using the esinet package for python. The open source code of the package can be found here: https://github.com/LukeTheHecker/invert.

## 7 Acknowledgements

We would like to thank Nvidia for sponsoring this work with a Titan V graphical computing unit as part of their academic seeding program. Furthermore, we thank all the contributors of the public data set as reported by Mikulan et al. (2020).

## 8 Appendix A

### 8.1 Forward Pass in the Fully Connected Neural Network

The full forward pass of the Fully Connected Neural Network is described as follows:

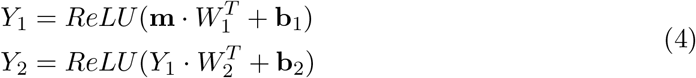

where:

*M ∈* ℝ ^*t×q*^ = EEG matrix

m *∈* R^*q*^ = EEG input at single time point

*k* = 300 = Number of neurons in hidden layers

*q* = 61 = Number of EEG channels

*p* = 1284 = Number of dipoles in source model

The output layer contains a linear activation function.

### 8.2 Forward Pass in the LSTM Network

Each LSTM cell contains three separate gates, namely the forget gate (*f*), the input gate (*i*) and an output gate (*o*) (Hochreiter & Schmidhuber, 1997). Each gate can be viewed as its own neural network with its weight matrices *U* and *W*. *σ* denotes the sigmoid function and tanh the hyperbolic tangent function. The candidate values for the new cell state are denoted in 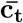.

The output of each gate is defined as follows:

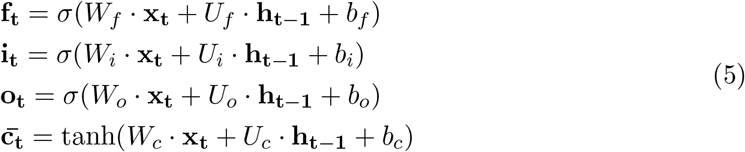

Where *X*_*t*_ is the input vector at time point *t* and **h**_*t−*1_ is the cell’s hidden state from the previous time point. *W* and *U* are weight matrices that are optimized during the training. Bias vectors are denoted by *b*.

The hidden state and cell state are updated as follows:

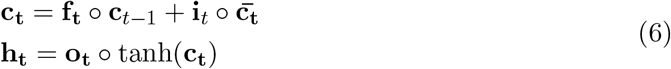

The ◦ operator denotes the Hadamard (i.e., element-wise) product.

Depending on the design of the LSTM model different variables may serve as the output of the LSTM cell. In our case, the hidden state at each time point was used. For seq2one models it is common to use only the final hidden state as output. In special cases, the cell state can be used as output, too.

The output layer of the LSTM model is a single linear fully-connected layer, through which the temporal outputs of the LSTM layers are passed sequentially.

Note, that in the present work, bidirectional LSTM cells were used. A bidirectional LSTM cell consists of two LSTM cells, one of which receives the time steps in reverse orders. This makes it possible to capture both past and future time points and thus improve predictions.

## 9 Funding

The project was funded by the European campus (eucor) seeding money.

## 10 Author Contribution

L.H., R.R. and J.K. designed the algorithm. M.M. evaluated the algorithm on real data. L.H, M.M. and J.K. and L.T.v.E. wrote the main manuscript. All authors reviewed the manuscript.

## 11 Conflict of Interest

The authors declare that there is no conflict of interest regarding the publication of this article.

## 12 Data Availability

All simulated data and evaluations can be recreated using the code which we will publish upon acceptance of this article. The SPES data used in this publication were published by Mikulan et al. (2020).

## Footnotes

1 http://surfer.nmr.mgh.harvard.edu/

2 https://github.com/lukethehecker/invert

## Notes

### Competing Interest Statement

The authors have declared no competing interest.

### Summary of Updates

Fixed inconsistent affiliations

## References

Abadi, M., Agarwal, A., Barham, P., Brevdo, E., Chen, Z., Citro, C., … Devin, M. (2016). Tensorflow: Large-scale machine learning on heterogeneous distributed systems. arXiv preprint arXiv:1603.04467.

Abeyratne, U. R., Zhang, G., & Saratchandran, P. (2001). EEG source localization: A comparative study of classical and neural network methods. International journal of neural systems, 11 (04), 349–359.

Awan, F. G., Saleem, O., & Kiran, A. (2019). Recent trends and advances in solving the inverse problem for EEG source localization. Inverse Problems in Science and Engineering, 27 (11), 1521–1536.

Baron, D., Duarte, M. F., Wakin, M. B., Sarvotham, S., & Baraniuk, R. G. (2009). Distributed compressive sensing. arXiv preprint arXiv:0901.3403.

Breakspear, M. (2017, March). Dynamic models of large-scale brain activity. Nature Neuroscience, 20 (3), 340–352. doi: 10.1038/nn.4497

Castanõ-Candamil, S., Höhne, J., Mart’ınez-Vargas, J.-D., An, X.-W., Castellanos-Dom’ınguez, G., & Haufe, S. (2015). Solving the EEG inverse problem based on space–time–frequency structured sparsity constraints. NeuroImage, 118, 598–612.

Chollet, F., et al. (2015). Keras.

Cui, S., Duan, L., Gong, B., Qiao, Y., Xu, F., Chen, J., & Wang, C. (2019). EEG source localization using spatio-temporal neural network. China Communications, 16 (7), 131–143.

Dale, A. M., Fischl, B., & Sereno, M. I. (1999). Cortical surface-based analysis: I. Segmentation and surface reconstruction. Neuroimage, 9 (2), 179–194.

Dale, A. M., & Sereno, M. I. (1993). Improved localizadon of cortical activity by combining EEG and MEG with MRI cortical surface reconstruction: A linear approach. Journal of cognitive neuroscience, 5 (2), 162–176.

de Peralta Menendez, R. G., Andino, S. G., Lantz, G., Michel, C. M., & Landis, T. (2001). Noninvasive localization of electromagnetic epileptic activity. I. Method descriptions and simulations. Brain topography, 14 (2), 131–137.

Destexhe, A., Contreras, D., & Steriade, M. (1999, June). Spatiotemporal Analysis of Local Field Potentials and Unit Discharges in Cat Cerebral Cortex during Natural Wake and Sleep States. The Journal of Neuroscience, 19 (11), 4595–4608. doi: 10.1523/JNEUROSCI.19-11-04595.1999

Dinh, C., Samuelsson, J. G., Hunold, A., & Hämäläinen, M. S. (2019). Contextual Minimum-Norm Estimates (CMNE): A Deep Learning Method for Source Estimation in Neuronal Networks., 14.

Dinh, C., Samuelsson, J. G., Hunold, A., Hämäläinen, M. S., & Khan, S. (2021). Contextual MEG and EEG source estimates using spatiotemporal LSTM networks. Frontiers in neuroscience, 15, 552666.

Fedorov, M., Koshev, N., & Dylov, D. V. (2020). Deep Learning for Non-invasive Cortical Potential Imaging. In Machine Learning in Clinical Neuroimaging and Radiogenomics in Neuro-oncology: Third International Workshop, MLCN 2020, and Second International Workshop, RNO-AI 2020, Held in Conjunction with MICCAI 2020, Lima, Peru, October 4–8, 2020, Proceedings (Vol. 12449, p. 45). Springer Nature.

Fischl, B., Sereno, M. I., Tootell, R. B., & Dale, A. M. (1999). High-resolution intersubject averaging and a coordinate system for the cortical surface. Human Brain Mapping, 8 (4), 272–284. doi: 10.1002/(sici)1097-0193(1999)8:4(272::aid-hbm10)3.0.co;2-4

Fuchs, M., Kastner, J., Wagner, M., Hawes, S., & Ebersole, J. S. (2002). A standardized boundary element method volume conductor model. Clinical Neurophysiology, 113 (5), 702–712.

Gavit, L., Baillet, S., Mangin, J.-F., Pescatore, J., & Garnero, L. (2001). A multiresolution framework to MEG/EEG source imaging. IEEE transactions on biomedical engineering, 48 (10), 1080–1087.

Godard, C., Matzen, K., & Uyttendaele, M. (2018). Deep Burst Denoising. In V. Ferrari, M. Hebert, C. Sminchisescu, & Y. Weiss (Eds.), Computer Vision – ECCV 2018 (Vol. 11219, pp. 560–577). Cham: Springer International Publishing. doi: 10.1007/978-3-030-01267-033

Gramfort, A., Kowalski, M., & Hämäläinen, M. (2012). Mixed-norm estimates for the M/EEG inverse problem using accelerated gradient methods. Physics in Medicine & Biology, 57 (7), 1937.

Gramfort, A., Luessi, M., Larson, E., Engemann, D. A., Strohmeier, D., Brodbeck, C., … Parkkonen, L. (2013). MEG and EEG data analysis with MNE-Python. Frontiers in neuroscience, 7, 267.

Gramfort, A., Strohmeier, D., Haueisen, J., Hämäläinen, M. S., & Kowalski, M. (2013, April). Time-frequency mixed-norm estimates: Sparse M/EEG imaging with nonstationary source activations. NeuroImage, 70, 410–422. doi: 10.1016/j.neuroimage.2012.12.051

Grech, R., Cassar, T., Muscat, J., Camilleri, K. P., Fabri, S. G., Zervakis, M., … Vanrumste, B. (2008). Review on solving the inverse problem in EEG source analysis. Journal of neuroengineering and rehabilitation, 5 (1), 25.

Grova, C., Daunizeau, J., Lina, J.-M., Bénar, C. G., Benali, H., & Gotman, J. (2006). Evaluation of EEG localization methods using realistic simulations of interictal spikes. Neuroimage, 29 (3), 734–753.

Hämäläinen, M. S., & Ilmoniemi, R. J. (1994). Interpreting magnetic fields of the brain: Minimum norm estimates. Medical & biological engineering & computing, 32 (1), 35–42.

He, B., Sohrabpour, A., Brown, E., & Liu, Z. (2018). Electrophysiological Source Imaging: A Noninvasive Window to Brain Dynamics. Annual Review of Biomedical Engineering, 20 (1), 171–196. doi: 10.1146/annurev-bioeng-062117-120853

He, K., Zhang, X., Ren, S., & Sun, J. (2016). Deep residual learning for image recognition. In Proceedings of the IEEE conference on computer vision and pattern recognition (pp. 770–778).

Hecker, L., Rupprecht, R., van Elst, L. T., & Kornmeier, J. (2020). ConvDip: A convolutional neural network for better M/EEG Source Imaging. bioRxiv.

Hochreiter, S., & Schmidhuber, J. (1997). Long short-term memory. Neural computation, 9 (8), 1735–1780.

Huang, G., Yu, Z. L., Wu, W., Liu, K., Gu, Z., Qi, F., … Liang, J. (2020). Electromagnetic Source Imaging via a Data-Synthesis-Based Denoising Autoencoder. arXiv preprint arXiv:2010.12876.

Ioannides, A. A., Bolton, J. P. R., & Clarke, C. J. S. (1990). Continuous probabilistic solutions to the biomagnetic inverse problem. Inverse Problems, 6 (4), 523.

Jiao, M., Wan, G., Guo, Y., Wang, D., Liu, H., Xiang, J., & Liu, F. (2022). A Graph Fourier Transform Based Bidirectional Long Short-Term Memory Neural Network for Electrophysiological Source Imaging. Frontiers in Neuroscience, 16.

Jin, K. H., McCann, M. T., Froustey, E., & Unser, M. (2017). Deep convolutional neural network for inverse problems in imaging. IEEE Transactions on Image Processing, 26 (9), 4509–4522.

Kingma, D. P., & Ba, J. (2014). Adam: A method for stochastic optimization. arXiv preprint arXiv:1412.6980.

Kornmeier, J., Friedel, E., Hecker, L., Schmidt, S., & Wittmann, M. (2019). What happens in the brain of meditators when perception changes but not the stimulus? PLoS One, 14 (10), e0223843.

Krizhevsky, A., Sutskever, I., & Hinton, G. E. (2012). Imagenet classification with deep convolutional neural networks. In Advances in neural information processing systems (pp. 1097–1105).

Leopold, D. A., Murayama, Y., & Logothetis, N. K. (2003, April). Very Slow Activity Fluctuations in Monkey Visual Cortex: Implications for Functional Brain Imaging. Cerebral Cortex, 13 (4), 422–433. doi: 10.1093/cercor/13.4.422

Mäkelä, N., Stenroos, M., Sarvas, J., & Ilmoniemi, R. J. (2018). Truncated rap-music (trap-music) for MEG and EEG source localization. NeuroImage, 167, 73–83.

Matsumoto, R., Kunieda, T., & Nair, D. (2017). Single pulse electrical stimulation to probe functional and pathological connectivity in epilepsy. Seizure, 44, 27–36.

Matsumoto, R., Nair, D. R., LaPresto, E., Najm, I., Bingaman, W., Shibasaki, H., & Lüders, H. O. (2004). Functional connectivity in the human language system: A cortico-cortical evoked potential study. Brain, 127 (10), 2316–2330.

McCulloch, W. S., & Pitts, W. (1943, December). A logical calculus of the ideas immanent in nervous activity. The bulletin of mathematical biophysics, 5 (4), 115–133. doi: 10.1007/BF02478259

Mégevand, P., Hamid, L., Dümpelmann, M., & Heers, M. (2019). New horizons in clinical electric source imaging. Zeitschrift für Epileptologie, 32 (3), 187–193.

Michel, C. M., & Brunet, D. (2019). EEG Source Imaging: A Practical Review of the Analysis Steps. Frontiers in Neurology, 10. doi: 10.3389/fneur.2019.00325

Michel, C. M., & Koenig, T. (2018, October). EEG microstates as a tool for studying the temporal dynamics of whole-brain neuronal networks: A review. NeuroImage, 180, 577–593. doi: 10.1016/j.neuroimage.2017.11.062

Mikulan, E., Russo, S., Parmigiani, S., Sarasso, S., Zauli, F. M., Rubino, A., … Pigorini, A. (2020, April). Simultaneous human intracerebral stimulation and HD-EEG, groundtruth for source localization methods. Scientific Data, 7 (1), 127. doi: 10.1038/s41597-020-0467-x

Nunez, P. L., & Srinivasan, R. (2006). Electric fields of the brain: The neurophysics of EEG. Oxford University Press, USA.

Pantazis, D., & Adler, A. (2021). MEG Source Localization Via Deep Learning. Sensors, 21 (13), 4278.

Pascual-Marqui, R., Michel, C. M., & Lehmann, D. (1994). Low-resolution electromagnetic tomography–a new method for localizing electrical activity in the brain. International Journal of psychophysiology, 18, 49–65.

Pascual-Marqui, R. D. (1999). Review of methods for solving the EEG inverse problem. International journal of bioelectromagnetism, 1 (1), 75–86.

Pascual-Marqui, R. D. (2007). Discrete, 3D distributed, linear imaging methods of electric neuronal activity. Part 1: Exact, zero error localization. arXiv preprint arXiv:0710.3341.

Razorenova, A., Yavich, N., Malovichko, M., Fedorov, M., Koshev, N., & Dylov, D. V. (2020). Deep Learning for Non-Invasive Cortical Potential Imaging. In Machine Learning in Clinical Neuroimaging and Radiogenomics in Neuro-oncology (pp. 45–55). Springer.

Robert, C., Gaudy, J.-F., & Limoge, A. (2002). Electroencephalogram processing using neural networks. Clinical Neurophysiology, 113 (5), 694–701.

Rosenblatt, F. (1958). The perceptron: A probabilistic model for information storage and organization in the brain. Psychological review, 65 (6), 386.

Schirrmeister, R. T., Springenberg, J. T., Fiederer, L. D. J., Glasstetter, M., Eggensperger, K., Tangermann, M., … Ball, T. (2017). Deep learning with convolutional neural networks for EEG decoding and visualization. Human brain mapping, 38 (11), 5391–5420.

Schuster, M., & Paliwal, K. K. (1997). Bidirectional recurrent neural networks. IEEE transactions on Signal Processing, 45 (11), 2673–2681.

Sun, R., Sohrabpour, A., Worrell, G. A., & He, B. (2022). Deep neural networks constrained by neural mass models improve electrophysiological source imaging of spatiotemporal brain dynamics. Proceedings of the National Academy of Sciences of the United States of America, 119 (31), e2201128119.

Sun, R., Sohrabpour, A., Ye, S., & He, B. (2020, May). SIFNet: Electromagnetic Source Imaging Framework Using Deep Neural Networks (Preprint). Bioengineering. doi: 10.1101/2020.05.11.089185

Tankelevich, R. (2019, February). Inverse problem’s solution using deep learning: An EEG-based study of brain activity. Part 1 - rel. 1.0.

Tropp, J. A., Gilbert, A. C., & Strauss, M. J. (2005). Simultaneous sparse approximation via greedy pursuit. In Proceedings.(ICASSP’05). IEEE International Conference on Acoustics, Speech, and Signal Processing, 2005. (Vol. 5, pp. v–721). IEEE.

van Lutterveld, R., Houlihan, S. D., Pal, P., Sacchet, M. D., McFarlane-Blake, C., Patel, P. R., … Brewer, J. A. (2017, May). Source-space EEG neurofeedback links subjective experience with brain activity during effortless awareness meditation. NeuroImage, 151, 117–127. doi: 10.1016/j.neuroimage.2016.02.047

Van Veen, B. D., van Drongelen, W., Yuchtman, M., & Suzuki, A. (1997, September). Localization of brain electrical activity via linearly constrained minimum variance spatial filtering. IEEE transactions on bio-medical engineering, 44 (9), 867–880. doi: 10.1109/10.623056

Wei, C., Lou, K., Wang, Z., Zhao, M., Mantini, D., & Liu, Q. (2021). Edge sparse basis network: A deep learning framework for EEG source localization. In 2021 International Joint Conference on Neural Networks (IJCNN) (pp. 1–8). IEEE.

Zorzos, I., Kakkos, I., Ventouras, E. M., & Matsopoulos, G. K. (2021). Advances in Electrical Source Imaging: A Review of the Current Approaches, Applications and Challenges. Signals, 2 (3), 378–391.

